# Network-based metrics of resilience and ecological memory in lake ecosystems

**DOI:** 10.1101/810762

**Authors:** David I. Armstrong McKay, James G. Dyke, C. Patrick Doncaster, John A. Dearing, Rong Wang

## Abstract

Some ecosystems undergo abrupt transitions to a new regime after passing a tipping point in an exogenous stressor, for example lakes shifting from a clear to turbid ‘eutrophic’ state in response to nutrient-enrichment. Metrics-based resilience indicators have been developed as early warning signals of these shifts but have not always proved reliable indicators. Alternative approaches focus on changes in the composition and structure of an ecosystem, which can require long-term food-web observations that are typically beyond the scope of monitoring. Here we prototype a network-based algorithm for estimating ecosystem resilience, which reconstructs past ecological networks solely from palaeoecological abundance data. Resilience is estimated using local stability analysis, and eco-net energy: a neural network-based proxy for ‘ecological memory’. We test the algorithm on modelled (PCLake+) and empirical (lake Erhai) data. The metrics identify increasing diatom community instability during eutrophication in both cases, with eco-net energy revealing complex eco-memory dynamics. The concept of ecological memory opens a new dimension for understanding ecosystem resilience and regime shifts, with eco-memory potentially increasing ecosystem resilience by allowing past memorised eco-network states to be recovered after disruptions.

## Introduction

The potential for stressed ecosystems to tip abruptly into a new regime has led to a proliferation of metrics attempting to quantify ecosystem resilience [1–10]. Lake eutrophication is a well-studied example of such a regime shift, and often exhibits evidence of alternative stable states and hysteresis [11–13]. Eutrophication occurs when increasing nutrient loading triggers positive feedbacks involving lake anoxia and light reduction within the ecosystem, driving a rapid shift from clear to turbid water conditions [14,15]. Recovery to clear conditions often requires a reduction in nutrient levels far below the original threshold, indicating hysteresis [13], although this is not a characteristic of all lake systems [11]. Prior to tipping, the ecosystem experiences declining resilience, defined here as the weakening of negative (stabilising) feedbacks relative to positive (amplifying) feedbacks, resulting in greater sensitivity to small shocks and longer recovery times (i.e. ‘engineering’ resilience) [16,17]. Attempts to develop ecosystem resilience indicators in order to understand and monitor the drivers of regime shifts broadly fall into three approaches: time-series metrics, compositional analysis, and network-based analysis.

### Monitoring ecosystem resilience

Many resilience indicators test for ‘critical slowing down’: a slowing recovery rate from perturbations and increasing variability, which have been detected in various environmental time-series [4–10]. These time-series metrics have had inconsistent success, however, when applied as early-warning signals (EWS) of tipping points in freshwater [18] and other ecosystems [19–22]. A key methodological issue is the need for prior knowledge of one response variable that captures the dynamics of the whole system when analysed. Other limitations include possible false positives or negatives (where EWS indicate an impending transition which never occurs, or is absent prior to a known transition), and sensitivity to subjective time-series analysis parameter choices [7,23,24]. These problems are partially resolved by using multiple sensitivity-tested metrics as generic resilience indicators rather than as EWS of specific critical transitions [16,25]. Palaeorecords present additional problems, with variable temporal resolution from changing sediment accumulation rates, compaction, or dating uncertainty making robust time-series analysis challenging [26,27].

As an alternative to metrics-based EWS, analysis of ecosystem composition seeks to detect changes in community functional dynamics without having to select or understand the full trophic ecology of any one species. To this end, Doncaster et al. [28] quantified the compositional disorder of diatoms and chironomids from sediment cores in three Chinese lakes, where low disorder signifies highly-nested sequential compositions. Several decades prior to critical transitions in these lakes, the correlation of disorder with biodiversity becomes negative and strengthens towards the tipping point. Theory and simulations [28] suggest that nutrient loading had shifted the competitive balance from ‘weedy’ (weakly competitive, fast-replicating) towards ‘keystone’ (strongly competitive, slow-replicating) species. This correlative approach avoids issues of variable temporal resolution, but the link with ecosystem resilience is indirect and the competition dynamics remain hypothetical. Another recent composition-based method identified negative skewness in nodal degree (connectivity) within a network of interacting diatom species, in response to increasing nutrient input into Chinese lake ecosystems [29]. This trend towards strong interactions is compatible with keystone dominance rising with exogenous stress.

Network-based methods represent another whole-ecosystem approach, which allows the possibility of performing local stability analysis on reconstructions of lake food-webs. Kuiper et al. [30] showed that food-web data in the form of material flux descriptions can be used to reconstruct interaction strengths between different species, which correspond to the interaction coefficients of a Lotka-Volterra ecosystem model (Figure 1). The matrix of these coefficients is equivalent to a Jacobian matrix of the Lotka-Volterra equations when considered as a dynamical system, and so methods for determining the local stability of dynamical systems can also be applied to this interaction matrix. These systems are considered locally stable if the real part of the Jacobian’s eigenvalues remain negative (representing stabilising net-negative feedbacks) [30–32]. A critical transition occurs when an eigenvalue crosses the zero line (representing the emergence of destabilising net-positive feedbacks). Given food-web measurements, the local stability of the food web can be estimated from the intraspecific interaction strengths, and is closely related to the Jacobian’s dominant eigenvalue, λ_d_ (the largest absolute eigenvalue of the Jacobian matrix) [30,33–35]. The value of λ_d_ increases with increasing post-perturbation recovery times as a result of the weakening of negative relative to positive feedbacks. It thus functions as a proxy for weakening ecosystem resilience. An application of this approach to the PCLake model system enabled tracking of destabilising food-web reorganisations prior to eutrophication [30].

**Figure 1:**
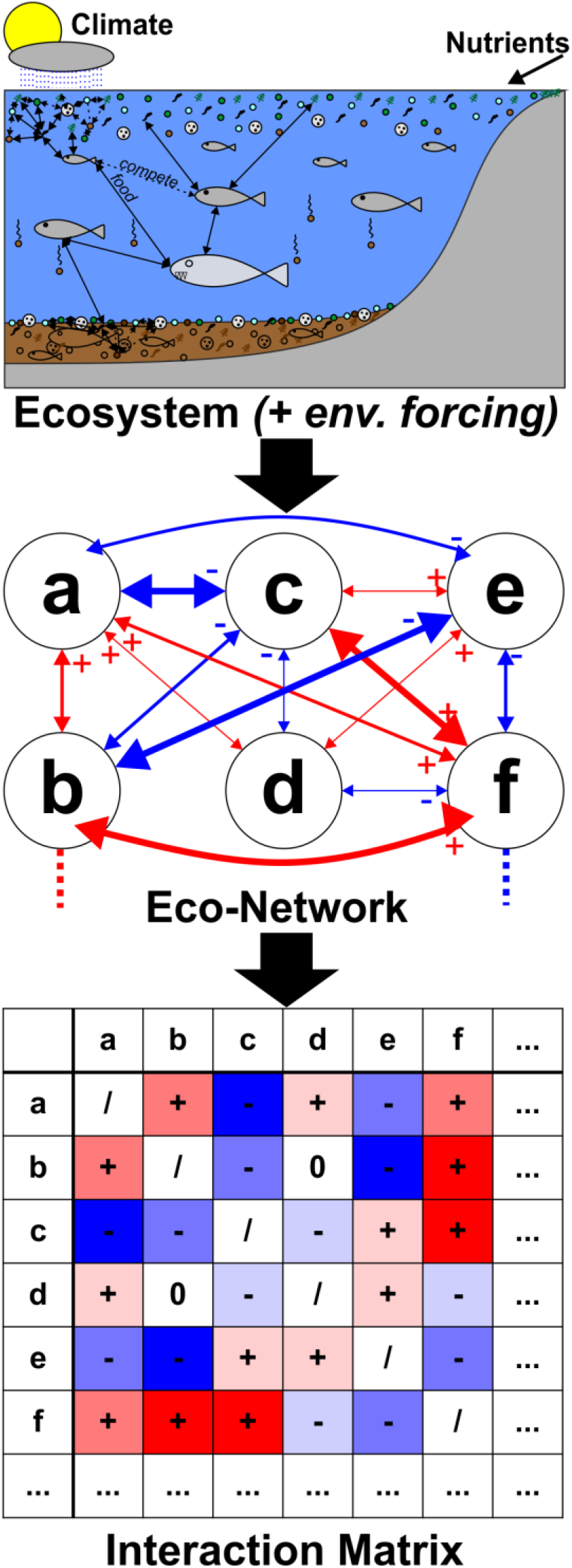
Schematic illustrating how ecosystem interactions (top) are represented as an eco-network (middle -lettered nodes represent individual species or functional groups, and edges their positive (red) or negative (blue) interactions), the structure of which forms an interaction matrix (bottom) suitable for resilience analysis. Only symmetric interactions are shown in this schematic; the model can also accommodate asymmetric interactions such as asymmetries in competitive (-/-) and mutualistic (+/+) interactions, and predator-prey (+/-) and host-parasite (-/+) interactions.

### Ecological network inference

Although the local network stability method depends less on temporal resolution than time-series analysis [30] and avoids reliance on choosing a single variable, it requires gathering detailed food-web data at regular intervals during eutrophication, a laborious [36] and often unmanageable task for real-world lakes. Many lakes nevertheless have palaeoecological records of species abundances through time, obtained from sediment cores. If past ecosystem interactions can be reconstructed from these data, it becomes possible to estimate past changes in ecosystem resilience. Network inference is a novel technique which allows ecological networks (eco-nets) to be reconstructed from abundance data [37–40]. This technique for characterising interspecific interactions comes from microbial metagenomics, where it provides an alternative to the unreliable proxy of assuming that correlations between the relative abundances of different species indicate causative interactions between them [39]. Network inference instead uses multilinear regression to infer the interaction coefficients of a discrete-time Lotka-Volterra ecosystem model from abundance data (detailed in Materials and Methods), with nodes in the inferred network representing individual species or functional groups and edges their interactions (Figure 1). Network inference thus allows the past structure of ecological communities to be reconstructed from palaeoecological abundance data, and is less sensitive to irregular temporal resolution than time-series analysis [38]. Ecosystem resilience can then be explored by performing local stability analysis on the inferred interaction matrix with a rolling temporal window, in order to track long-term changes in λ_d_.

### Ecological memory and eco-net energy

In this study we also develop and apply a novel neural network-based method of resilience analysis. Analyses of Lotka-Volterra systems demonstrate how an ecosystem can retain a distributed ‘memory’ of past states as a result of a process akin to unsupervised Hebbian learning in a type of neural network known as a Hopfield network [41,42]. Neural networks are computing systems loosely based on brains, with nodes (neurones) connected by edges (axons/synapses) used to process information. Neural networks are trained to recognise and process particular input data (the training input), with ‘learning’ indicating adaptation of the network’s connections in response to the training, and the training input then stored as a distributed ‘memory’ as the network’s structural adaptations (Figure 2a). In many networks this process is externally guided to reach desired results (‘supervised learning’) and is not applicable in natural ecosystems, but unsupervised ‘Hebbian’ learning can also lead to memory spontaneously emerging when some neural networks are given training input. In Hopfield networks for example, correlation in activity between two nodes leads to an increase in the connection strength between them – a process described colloquially as *“neurons that fire together, wire together”* (Hebb’s Rule [43–46]). Over time this allows the emergence of distributive associative memory of training input that can be recovered when given degraded input (Figure 2b).

**Figure 2:**
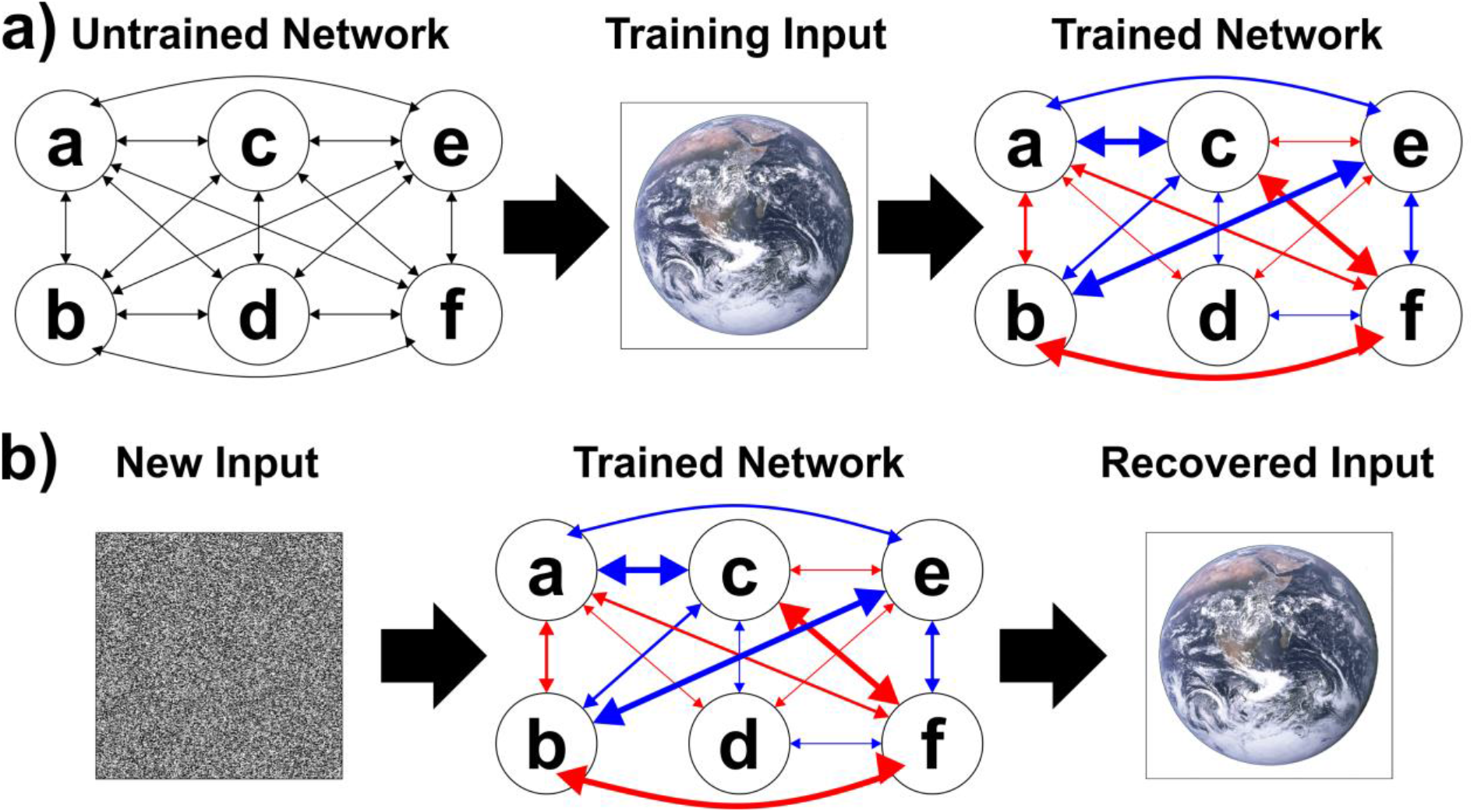
Schematic illustrating how (a) neural (or ecological) networks are trained by exogenous forcing (with environmental drivers), and (b) the character of this training input can be recovered after applying new degraded input to the trained network instead. In this case relearning is not enabled and so the new input does not lead to network alterations. Lettered nodes represent individual species or functional groups, and edges their positive (red) or negative (blue) interactions; black arrows in the untrained network indicate initial null or random values prior to training. Only symmetric interactions are shown in this schematic; the model can also accommodate asymmetric interactions such as asymmetries in competitive (-/-) and mutualistic (+/+) interactions, and predator-prey (+/-) and host-parasite (-/+) interactions.

Power et al. [41] proposed that eco-nets experience similar learning dynamics, with species (nodes) that frequently co-occur developing stronger interactions (edges), allowing emergence of a distributed associative “ecological memory” (eco-memory) of past environmental forcing (training input) that acts as a stable attractor (Figure 2). This process has been simulated in a simple Lotka-Volterra model in which species are subject to natural selection by applying environmental forcing in the form of altered individual carrying capacities (equivalent to training input), with this training input subsequently recoverable even when the model is reset with new abundances [41]. This ecological memory can increase the ecosystem’s resilience by acting as a stable attractor towards which the ecosystem recovers after perturbation. These findings suggest that ecosystems can learn from environmental forcing and become more resilient as a result of network effects [42], but this process has not yet been tested in more realistic process-based models or empirical data.

In this paper, we quantify the strength of ecological memory from abundance data as a metric of resilience, rather than recovery of training inputs *per se.* Although memory strength has no direct metric, one can calculate a Hopfield network’s ‘energy’. Training a Hopfield network results in a stable minimum forming in its associated energy function, which represents the memorised state. Once trained, this minimum acts as an attractor when the network is given new input (Figure 3a). Because network energy decreases with learning, and is minimised at locally stable points, it indicates how much the network has learnt about its past training. We can calculate eco-net energy, E_N_, by treating the interaction matrix reconstructed by our network-inference algorithm as the weight matrix [41] of a continuous Hopfield network [43–46] (see Materials and Methods), and use it to indicate the proximity of the eco-net to the bottom of a memorised attractor as well as the depth of the attractor (Figure 3, right). We would expect low E_N_ for eco-nets that have ‘learnt’ from stable environmental conditions or have shifted to a more stable memory attractor (Figure 3a), and higher E_N_ when ecosystem destabilisation perturbs the eco-net away from its learned state or shifts it to a less stable memory attractor (Figure 3b). Unlike a conventional Hopfield network though, eco-nets can also continue to learn from new conditions. Thus, a novel stable ecosystem state can form a new memory (‘relearning’) and energy minimum, potentially leading to lower E_N_ after some initial destabilisation (Figure 3c).

**Figure 3:**
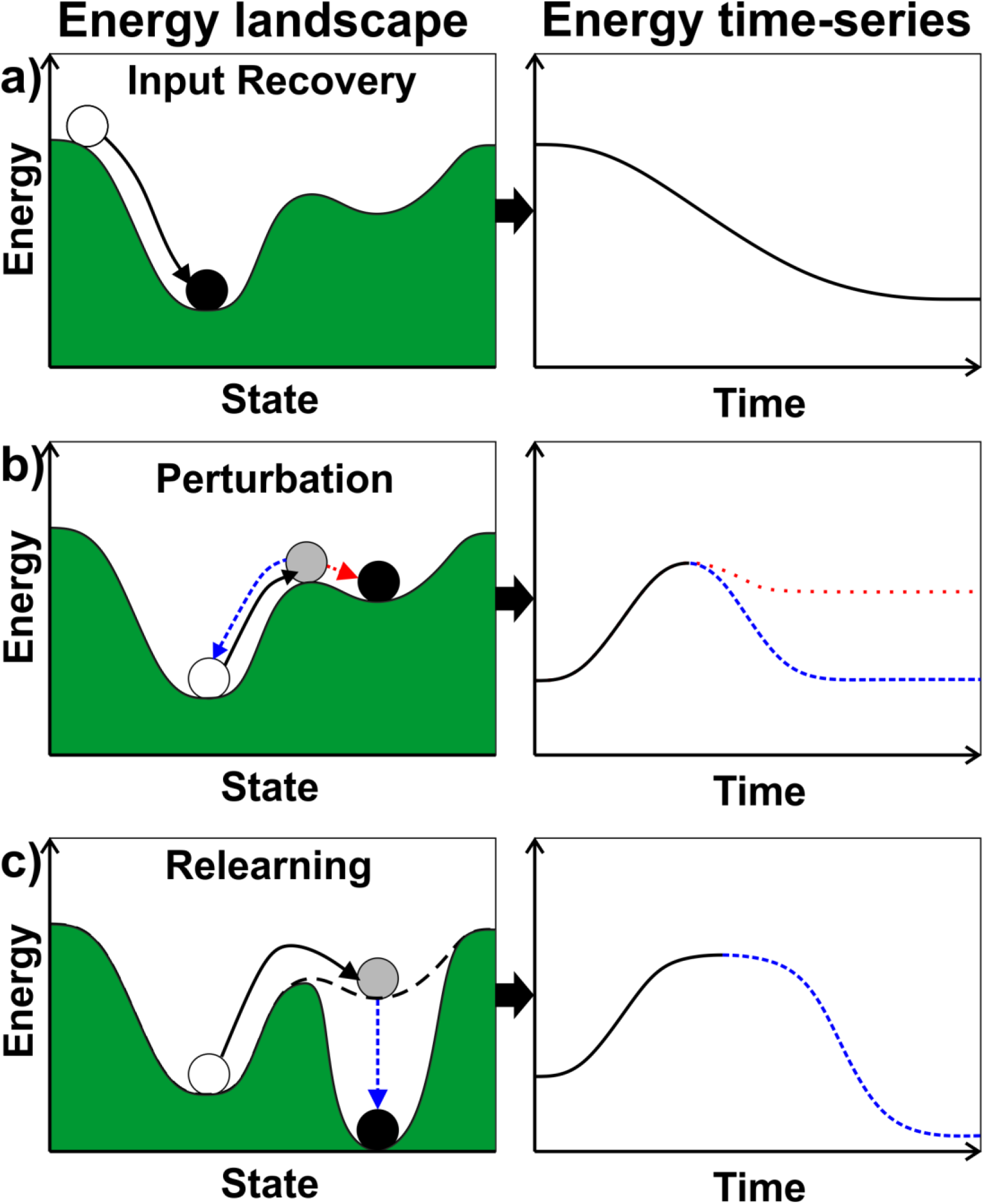
Schematic illustrating the dynamics of network energy in a neural network with attractor “memories”. In (a) the neural network has formed an energy landscape (left) after learning from training input, allowing the stored memory to be recovered if the input is reset, as illustrated by the energy of the ball declining (right) as it moves into the energy minima (i.e. the memorised state). In (b), the state of the network is then perturbed (left) leading to a temporary increase in the network’s energy (right), followed by the network either returning to the original attractor and low energy state (dashed blue arrow/line) or passing into an alternative, less stable attractor with higher energy (i.e. a different, weaker memory – dotted red arrow/line), In (c) network relearning is possible, and the energy landscape adjusts to allow the alternative attractor/memory to deepen in the network’s new state (left, dashed blue arrow), leading to the network eventually reaching an even lower energy than before (right, dashed blue line).

### Study Aims and Design

In this paper we prototype and test an algorithm in R [47] for reconstructing ecological networks (eco-nets) from palaeoecological data and determining past changes in resilience using network-based metrics. We first use network inference to reconstruct the ecosystem’s Lotka-Volterra interaction matrix from abundance data on a rolling temporal window, before exploring the ecosystem’s changing resilience using two different approaches. Firstly, we perform local stability analysis on the inferred interaction matrices in order to track changes in λ_d_. We test this analysis on two different datasets of eutrophication-induced regime shifts: whole ecosystem output from a commonly-used lake ecosystem model (PCLake+) for a hypothetical lake, and empirical community-level data from lake Erhai in China, where a critical transition was observed in 2001 [48]. Secondly, we calculate the neural network energy E_N_ on the same data for both test-cases in order to evaluate our theoretical expectations for E_N_. For comparison with conventional leading indicators of critical transitions, we also calculate time-series metrics (TSMs: autoregressive coefficient at lag 1 (AR1), standard deviation (SD), skewness, and kurtosis), biodiversity, and for empirical data the sequential disorder-biodiversity correlation. Detailed methods and scripts are available in the Materials and Methods section. Both metrics find evidence of increasing diatom community instability during eutrophication in each dataset, with eco-net energy revealing complex eco-memory dynamics occurring before and during the regime shift.

## Results and Discussion

### Model data: PCLake+

We first apply the algorithm to output from a default setup of PCLake+ [49], an extension of the widely-used PCLake model of lake eutrophication, as a test-bed with well-known dynamics and drivers for generating realistic data. The artificial lake supports 14 functional groups of species, and has phosphorus input increasing along a nonlinear but non-hysteretic load-response curve to induce eutrophication (detailed in Materials and Methods). The impact of nutrient enrichment by phosphorus is clearly visible across three distinct phases in lake conditions, dominant eigenvalue (λ_d_; i.e. local instability), eco-net energy (E_N_), and biodiversity (Figure 4a-d, left). During Phase 1, initial destabilisation, λ_d_ and E_N_ both increase after the start of nutrient input and the replacement of vegetation by plankton, with both peaking sharply at the phase’s end along with biodiversity. At the beginning of Phase 2, pretransition, declining zooplankton abundance is marked by falls in both λ_d_ and E_N_, followed by both plateauing with the continued rise in phytoplankton (indicated by chlorophyll concentration). Phase 3, regime shift, is marked by initial peaks followed by steady declines in both λ_d_ and E_N_, with biodiversity continuing to decline towards eutrophic conditions.

**Figure 4:**
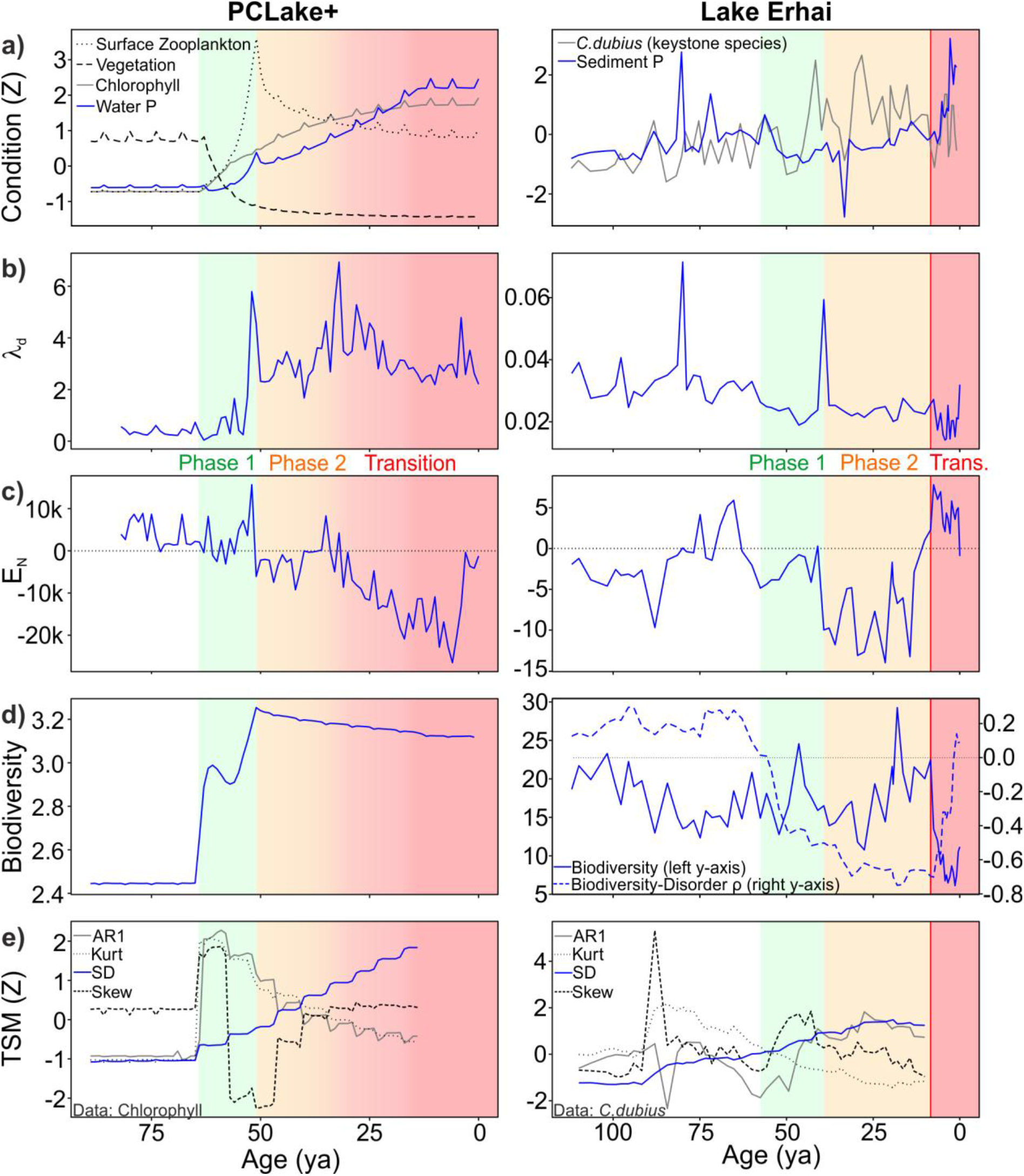
Time-series of lake conditions and resilience metrics for PCLake+ (left) and lake Erhai (right) through time (years ago) under increasing concentrations of phosphorus (P) leading to eutrophication. (a) Lake conditions indicated by normalised concentrations/abundances of vegetation, zooplankton, and chlorophyll (phytoplankton) in PCLake+, and increased keystone species (*Cyclostephanos dubius*) abundance in Erhai; (b) dominant eigenvalue (i.e. local instability) λ_d_; (c) eco-net energy E_N_; (d) biodiversity (inverse Simpson index), and disorder-biodiversity correlation (Erhai only, [28]); (e) normalised time-series metrics (TSM; for comparison only). Phase 1: initial destabilisation (green); Phase 2: pre-transition (orange); Phase 3: regime shift (red, red vertical line indicates known critical transition).

These trends in λ_d_ and E_N_, and the fluctuations around them, indicate complex eco-memory dynamics starting decades before the transition. The initial destabilisation away from memorised conditions, reflected in the rising λ_d_ indicative of weakened net-negative feedbacks, was followed by multi-phase “relearning” processes, reflected in decreasing E_N_, as the eco-net reconfigured its dominant functional groups. Because PCLake+ was initialised and run as a stable non-eutrophic system prior to exogenous forcing by nutrient input, the system initially had only this one memorised attractor in its energy landscape. The observation of declining E_N_ after an initial destabilisation phase indicates learning of a new and more stable attractor (as conceptualised in Figure 3c). Increasing the sediment accumulation rate through the artificial core does not alter the overall signal but reduces the temporal resolution of deeper signals (Supplementary Figure S1), indicating this methodology is not overly sensitive to temporal resolution.

### Empirical data: lake Erhai

Similar to our PCLake+ scenario, lake Erhai in south-western China transitioned abruptly to sustained eutrophication after decades of exogenous forcing from phosphorus input, which were largely absorbed by lake sediment prior to the transition. Diatom abundances were sampled at regular radioisotope-dated intervals down sediment cores [28,48]. Because they are well preserved through time, the time-series of diatom composition provides a useful indicator of ecosystem state for producer-phytoplankton functional groups. We posit that the community-level λ_d_ for diatoms acts as a proxy for whole-ecosystem resilience, given that diatoms play a key role in the trophic loops involved in eutrophication [30] and are ecologically sensitive to water quality [50,51]. Despite not having data on the whole ecosystem network, in common with almost all empirical studies, we can still expect λ_d_ and E_N_ to record changes in wider network connections during a major regime shift that impacts on the whole ecosystem.

We observe comparable phases of activity to PCLake+ (Figure 4a-d, right). Phase 1, initial destabilisation, begins with the start of nutrient enrichment ~50-60 ya, as reported in historical records [48]. This phase is marked by an increasingly negative biodiversity-disorder correlation in diatom composition, indicative of strengthening interactions between species [27], accompanied by rising E_N_ towards a brief peak in λ_d_. In Phase 2, pre-transition, λ_d_ sustains a low plateau and E_N_ drops to a low fluctuating average, while the biodiversity-disorder correlation continues its increasingly negative trend. Just prior to the observed transition at ~8 ya, E_N_ increases rapidly to its highest level. In Phase 3, regime shift, both E_N_ and λ_d_ drop slightly, accompanying the sharp drop in biodiversity and the return to positive disorder-biodiversity correlation indicative of weakly interacting species [27].

These results show a similar pattern to the PCLake+ results. Phase 1 is marked by increasing E_N_ and a sharp peak in λ_d_ indicative of the eco-net destabilising decades before the transition. At the start of Phase 2, the sharp drop in E_N_ indicates relearning of newly forming eco-net configurations. Lower E_N_ could also represent the network shifting to a pre-existing alternative and deeper memory attractor, as lake Erhai may have experienced and memorised eutrophic conditions before (the reverse of Figure 3b). The subsequent rapid increase in E_N_ up to the transition into Phase 3 is consistent with a composition dominated by strongly interacting species for which the system has no previous memory. The post-transition decline in λ_d_ and E_N_ partly mirror PCLake+ but the decline is smaller for E_N_, which may result from slower forcing (Supplementary Figure S2) or the full decline has yet to be observed. The overall similarity in phasing between lake Erhai and PCLake+ suggests that diatoms can act as a community-level proxy for the whole lake ecosystem. However, the progression of λ_d_ and E_N_ prior to Phase 1 also shows marked changes in magnitude, suggesting that these metrics respond also to events unrelated to the eventual regime shift.

### Comparison to time-series metrics

TSMs are shown for comparison as a conventional resilience indicator method. As a consistently sampled model lake with constant temporal resolution, PCLake+ provides idealised conditions for observing TSMs during eutrophication (Figure 4e). Prior to a critical transition autocorrelation (AR1) and variability (SD) are expected to increase as a result of ‘critical slowing down’, while kurtosis may also increase and skewness may change either way [6]. Phase 1 has rapidly peaking AR1, skewness, and kurtosis, while SD begins a steady increase. Phase 2 has declines in all metrics, except for SD which continues its steady rise. Lake Erhai shows the largest changes in metric magnitudes prior to Phase 1 at ~90 ya, which may represent an unrecorded precursor event. Its Phase 1 nevertheless has rising AR1, skewness and SD as seen in PCLake+, but a steady decline in kurtosis (in contrast to PCLake+). Its Phase 2 has declines in skewness and kurtosis while SD continues to increase (consistent with PCLake+), but AR1 stays high (in contrast to PCLake+). Both examples suggest SD consistently increases during destabilisation, but variable temporal resolution can obscure this in real-world data (Supplementary Figure S1). However, due to the methodological limitations described earlier, TSM are not considered robust for lake Erhai without further sensitivity and significance testing, and are shown for comparison only.

### Further Development

Future iterations of these metrics for resilience and ecological memory should aim to address some current limitations. The network-inference algorithm functions most efficiently on long datasets that sufficiently resolve pre-perturbation equilibrium population dynamics and had sensitive detection thresholds to minimise data sparseness, but few palaeorecords meet this ideal. Recent developments in metagenomic network inference [38] suggest scope for improving efficiency on short or sparse data. We have assumed that ecosystem dynamics follow a generalised Lotka-Volterra model, in which all abundance changes are caused by linear pairwise interspecific interactions, and all other processes are pooled into a noise term. However, nonlinear interactions are expected in some ecosystems [37,52–56] and potentially allow a better representation of multiple alternative stable states [57,58], but are harder to parameterise. Future work will assess the feasibility of allowing nonlinear functional responses. The model currently includes only biotic interactions, with life-environment interactions implicit in parameter constants. Incorporating life-environment interactions explicitly would allow for more realistic feedback loops to emerge (such as anoxia-driven phosphorus release from sediment) that are critical for understanding resilience. However, the eco-net cannot simply be extended to include abiotic elements, because the environment is assumed to be its training input. Possible solutions include neural networks that allow training input that is dynamic (e.g. continuous-time recurrent neural networks) and interacts with the eco-net itself (e.g. multi-layer or adversarial networks).

### Conclusions

Our analyses indicate that changes in past ecosystem resilience and memory can be reconstructed in palaeoecological datasets of community composition from various settings, providing complementary information to metrics-based early warning signals. Our development here of eco-net energy has allowed us to explore eco-memory in process-based models and empirical data for the first time. Further work is required to fully understand the drivers and implications of eco-net energy dynamics, and to disentangle the effects of eco-memory from other drivers of ecosystem resilience. Eco-memory opens up a new dimension for understanding ecosystem resilience, with the formation of eco-memory potentially increasing resilience by allowing past stable eco-network states to be recovered after disruptions.

## Materials and Methods

### Ecological network inference

All analysis was conducted in R [47]. Abundance data-in the form of the relative proportional abundance per species at regularly sampled intervals from either empirical or model sources – were first transformed by isometric log ratio to account for compositionality in relative data and imputed to counter data sparseness [59] to form abundance matrix ***X***. To find the structure of the eco-net, we followed the methods of metagenomics network inference [39,40]. We first assumed that ecological interactions follow a generalised Lotka-Volterra model:

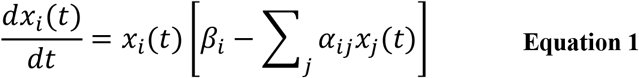

where *x* is the abundance of species *i* or *j, α_ij_* is the interaction coefficient between species *i* and *j* (<—1 < *α_ij_* < 1), and *β_t_* is the constant unitary net growth rate of species *i*. Taking the logarithm of a discrete-time Lotka-Volterra model based on Equation 1 and assuming constant time-steps [39] gives:

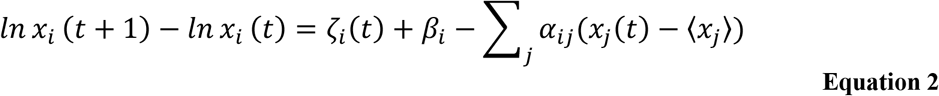

where *ζ* is the noise term and 〈*x_j_*〉 is the median abundance (approximating to equilibrium abundance and carrying capacity for normalising against; assumes sufficient data to capture equilibrium population dynamics and ephemeral species). Given abundance time-series, the interaction coefficients *α_ij_* can be found by performing multiple linear regression (lasso regularised due to data sparseness and limited observations relative to variables, and k-Fold cross-validated with at least 2 iterations to improve prediction performance) of [*lnx_i_* (*t* + 1) – *ln x_i_* (*t*)] against [*x_j_*(*t*) — 〈*x_j_*〉]. The regression coefficients then populate the elements of interaction matrix ***A***, which represents the eco-net’s structure at time *t*. The interaction matrix was calculated from the abundance data on a 50% rolling window in order to capture changes in the eco-network over time. We use non-interpolated data to avoid well-known biases interpolation introduces to time-series analysis [26], although this results in slightly varying time-steps. Network inference and linear regression are less sensitive to non-equidistant time points than time-series analysis [38], but the discrete-time Lotka-Volterra model still assumes constant time-steps. To minimise this limitation, input data should be scrutinised prior to analysis and sections with substantial shifts in time resolution excluded.

### Local Stability Analysis

The eco-net’s stability was estimated using local stability analysis, by taking the absolute of the interaction matrix ***A*** (which is equivalent to the Jacobian of the Lotka-Volterra system [30–32]; the system is considered stable at its equilibrium point if the real part of the Jacobian’s eigenvalues remain negative, indicating net negative feedbacks), setting intraspecific interaction (*α_ii_*) to 0 to satisfy the Perron-Frobenius Theorem [33], and calculating the dominant eigenvalue λ_d_ of the new positive matrix ***<u>A</u>***. The Jacobian’s λ_d_ is tightly related to other ecosystem stability metrics, having been shown to be equivalent to the smallest value for *α_ii_* necessary for the interaction matrix ***A*** to be quasi-diagonally dominant and therefore stable, as well as being related to the maximum loop weight of the longest trophic loop [30,33–35]. λ_d_ was calculated on the rolling interaction matrix in order to estimate changes in ecosystem stability over time.

### Ecological memory and eco-net energy

Recent theory suggests that ecosystems may have a distributed ‘memory’ of past states as a result of a process akin to unsupervised Hebbian learning in a Hopfield network [41,42] (see Introduction). There is no direct metric of Hopfield network memory strength, but we can calculate its energy, which is minimised at stable points. By assuming the eco-net’s interaction matrix ***A*** is equivalent to a neural network’s weight matrix ***W*** [41] and treating it as a continuous Hopfield network, we posit the eco-net energy at time *t*, E_N_, can be calculated by:

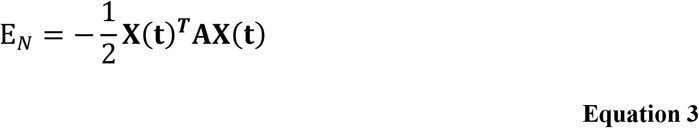

where ***X***(***t***) is a vector of species abundances at time *t* [43–46]. We expect low E_N_ for eco-nets that have ‘learnt’ from stable environmental conditions or have shifted to a more stable attractor (Figure 3a), and higher E_N_ when destabilisation shifts the eco-net away from its learned state or to a less stable attractor (Figure 3b). However, asymmetric matrices with nonzero diagonals, such as an ecosystem interaction matrix, can lead to chaotic behaviour and the network converging to alternative metastable points, meaning eco-nets may not always completely minimise their energy.

### Comparison Metrics

For comparison we also calculated normalised time-series metrics (TSMs: Z-scores of AR1, standard deviation (SD), skewness, and kurtosis) on the abundance of a previously identified keystone species *(Cyclostephanos dubius;* Erhai [28,48]) or surface chlorophyll concentration (PCLake+) with variance-stabilising square-root transformation, Gaussian detrending (bandwidth: PCLake+ 1%; Erhai 5%), and a rolling window of 50% up to but not including the transition (observed critical transition for lake Erhai, nutrient-enrichment ceasing for PCLake+) (R: *generic_ews, “earlywarnings”[6]).* However, due to the methodological limitations described in the Introduction [7,23,24,26,27], time-series metrics are shown for comparison only. We also calculate compositional disorder metrics [28] (disorder/nestedness, biodiversity (inverse Simpson index), and their correlation using pretransition window; R: *nestedtemp* & *diversity, “vegan”[60])* where possible.

### PCLake+ Modelling

PCLake+ is initialised in its default clear deep state with monthly time-steps, run for 200 years to ensure stability, and nutrient input increased from 0.0001 to 0.005 gPm^-2^d^-1^ (with the latter value considered sufficient to induce a eutrophic regime shift in PCLake [61,62]) over 50 years to induce eutrophication. Eutrophication is visible in the results as a shift in dominance from vegetation to plankton, which is non-abrupt as PCLake+ in its default settings displays a nonlinear but non-hysteretic load-response curve [49,63]. Model outputs consist of dry-weight abundances for each functional group (diatoms, green algae, blue-green algae, vegetation, zooplankton, and detritus split into surface and benthic groups, along with fish split into adult, juvenile, and piscavorous). As real-world palaeodata has reduced resolution compared to the daily/monthly model output as a result of sedimentary preservation, it was necessary to bin the model output into intervals equivalent to depth sampling in order to generate realistic artificial lake palaeorecords. This also allowed us to simulate the differing effects of constant or variable accumulation rates on our results (Supplementary Figure S1). The binned model outputs are then used as abundance inputs for the algorithm. We cannot compute nestedness or nestedness-biodiversity correlation for PCLake+ as only functional groups which always have some presence (rather than individual species which can go locally extinct) are available.

## Acknowledgements

Erhai data was provided by the Nanjing Institute of Geography and Limnology. We thank Pete Langdon and Richard Watson for comments on preliminary results, Annette Janssen for advice on using PCLake+.

