## Supplementary for "Network-based metrics of resilience and ecological memory in lake ecosystems"

### 1. Impact of increasing sediment accumulation rates on PCLake+ results

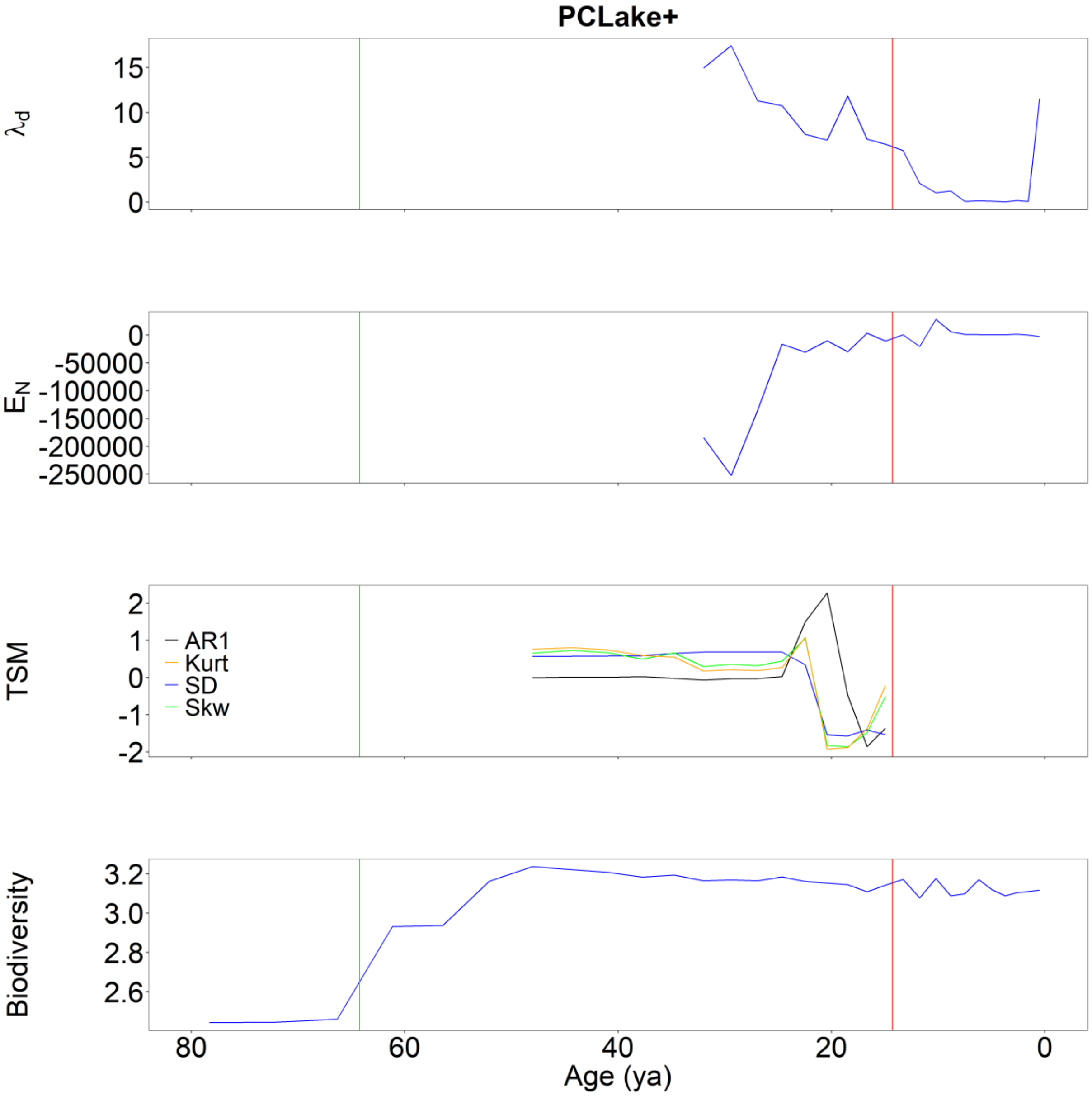


**Figure S1**: Results illustrating the impact of an increasing accumulation rate downwards through the core on PCLake+ resilience metrics (age-depth model provided in PCLake+ output file). This illustrates that broadly similar trends (elevated $\lambda_d$ declining towards transition, and low $E_N$ in phase 2 increasing across the transition as nutrient input stabilises) can still be seen as for the constant accumulation rate analysis, but their resolution is truncated and timings slightly offset, limiting the interpretation. In contrast, TSM are very different to the constant accumulation rate results, illustrating their higher sensitivity to temporal resolution. Vertical lines indicate nutrient-enrichment start (green) and finish (red).

#### 2. PCLake+ slower nutrient enrichment runs

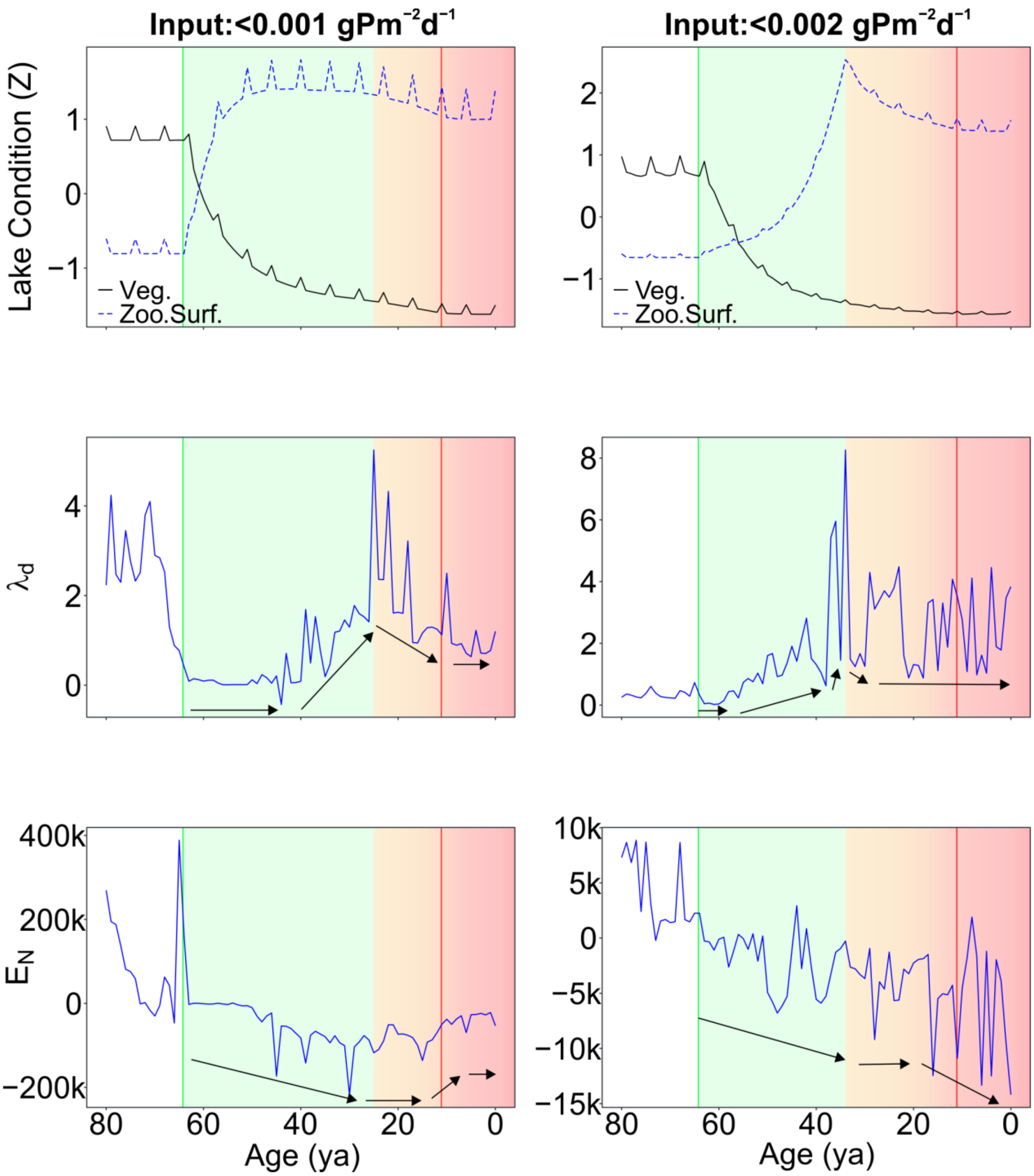


**Figure S2:** Impact of smaller increases in nutrient input on PCLake+ $\lambda_d$ and $E_N$ results for slower nutrient enrichment (slowest, left; moderate, right) than our main results. (Top) Lake conditions indicated by normalised concentrations/abundances of vegetation-zooplankton and chlorophyll; (middle) dominant eigenvalue (i.e. local instability) $\lambda_d$; (bottom) eco-net energy $E_N$. Slower nutrient enrichment results in a longer phase 1 with no end-phase $E_N$ peak, suggesting the ecosystem destabilisation indicated by $\lambda_d$ is slow enough for continuous eco-net relearning. Slower nutrient enrichment also allows for a post-transition increase in $E_N$ in the slowest scenario (left), unlike in the mid-range (right) or main text scenarios where $E_N$ decreases substantially after the transition, suggesting that further ecosystem reorganisation and relearning occurs after more intense perturbations. High values before phase 1 for the slowest scenario appear to be data artefacts. Phase 1: initial destabilisation (green); Phase 2: pre-transition (orange); Phase 3: regime shift (red); vertical lines indicate nutrient-enrichment start (green) and finish (red).
